# Linkage disequilibrium and population structure in a core collection of *Brassica napus* (L.)

**DOI:** 10.1101/2021.04.06.438572

**Authors:** Mukhlesur Rahman, Ahasanul Hoque, Jayanta Roy

**Author notes:** These authors have contributed equally to this work and share first authorship (Mukhlesur Rahman, Ahasanul Hoque).

## Abstract

Estimation of genetic diversity in rapeseed/canola is important for sustainable breeding program to provide an option for the development of new breeding lines. The objective of this study was to elucidate the patterns of genetic diversity within and among different structural groups, and measure the extent of linkage disequilibrium (LD) of 383 globally distributed rapeseed/canola germplasm using 8,502 single nucleotide polymorphism (SNP) markers. The germplasm accessions were divided into five subpopulations (P1 to P5) with obvious geographic and growth habit-related patterns. All subpopulations showed moderate genetic diversity (average *H* = 0.22 and *I* = 0.34). The pairwise *F_st_* comparison revealed a great degree of divergence (*F_st_* > 0.24) between most of the combinations. The rutabaga type showed highest divergence with spring and winter types. Higher divergence was also found between winter and spring types. Overall, mean linkage disequilibrium was 0.03 and it decayed to its half maximum within < 45 kb distance for whole genome. The LD decay was slower in C genome (< 93 kb), relative to the A genome (< 21 kb) which was confirmed by availability of larger haplotype blocks in C genome than A genome. To maximize genetic gain, hybridization between rutabaga and other types are potentially the best option. Hybridization between spring and winter, semi-winter type is also helpful to maximize the diversity in subsequent populations. Low genetic differentiation between both spring type subpopulations (P4 and P3) will accelerate favorable allele accumulation for specific traits in elite lines. The Neighbor-Joining tree and kinship matrix will assist to identify distantly related genotypes from subpopulations to utilize in hybridization. The low levels of LD and population structure make the core collection an important resource for association mapping efforts to identify genes useful in crop improvement as well as for selection of parents for hybrid breeding.

## Introduction

Rapeseed/canola (*Brassica napus L*., AACC, 2n = 4x = 38), is a recent allopolyploid of polyphyletic origin that evolved from hybridization events between two parental ancestors of *B. oleracea* (Mediterranean cabbage, CC, 2n = 2x = 18) and *B. rapa* (Asian cabbage or rapeseed, AA, 2n = 2x = 20) (1). Rapeseed/canola is the second largest oilseed crops produced in the world after soybean (2). Canola oil is mostly used in frying and baking, margarine, salad dressings, and many other products. Because of its fatty acid profile and the lowest amount of saturated fat among all other oils, it is commonly consumed all over the world and is considered a very healthy oil (3). Canola oil is also rich with alpha-linolenic acid (ALA), which is associated to a lower risk of cardiovascular disease (3). Additionally, canola is utilized as a livestock meal and is the second largest protein meal in the world after soybean (4). In the United States of America, the canola production increased 13.5 folds from five years average of 1991-1995 (0.11 m tons) to five years average of 2015-2019 (1.49 m tons) (5). Due to the growing importance of canola there is a constant need to improve its yield which can be negatively affected by biotic and abiotic stresses. Canola expresses three growth habits, winter, spring and semi-winter. The spring canola is planted in the early spring and harvested in the late spring of the same growing season (6). The winter type canola is seeded in the fall, vernalized over the winter to induce flower and harvested in the summer (6). The semi-winter type is needed for a shorter period of vernalization to induce flower (7). In order to adapt to different growing regions, plants developed systems that sense temperature, light quality, day length, as well as stress signals (8–11). Plants from colder zones require vernalization to flowering (12). Also day length and light quality affect the ability to flower after winter (13). The transition from the vegetative to flowering stage is also controlled by plant hormones and the circadian clock (14, 15). Multiple studies suggest the Arabidopsis flowering-time gene network might be similar to the one in Brassica (16, 17). Detailed understanding of the nature of genes involved in the growth habits of canola will facilitate development of cultivars adapted to different latitudes (18).

Due to limited use of diversified germplasm in breeding program, development of superior cultivars through traditional breeding might become unsuccessful and lead to stagnation in plant improvement (19). The recent origin of *B. napus* as a species and its very recent domestication (400 years ago), as well as selection on few phenotypes (e.g. low erucic and glucosinolate acids, seed yield) contributed to the low diversity which threatens sustainable production of the crop (20). The narrow genetic diversity might also limit the prospects for hybrid breeding where complementing genepools are needed for the optimal exploitation of heterosis (21). Therefore, it is crucial to study, preserve, and even introduce genetic diversity into rapeseed since the diversity is the best source of biotic and abiotic stress resistance, and various agronomical and morphological traits. Canola improvement can benefit from the availability and detailed characterization of genetically diverse germplasm. The knowledge of population structure, genetic relatedness, and patterns of linkage disequilibrium (LD) are also prime requirements for genome-wide association study (GWAS) and genome selection directed breeding strategies (22, 23).

Multiple genetic diversity and population structure studies, based on LD, have already provided information in regards to genetic diversity in various *B. napus* collections around the world (24–27). Unfortunately, there is a limited number of studies investigating the genetic variation of canola germplasms in the U.S. core collection justifying a great need for such a research focus. The LD analysis provides an important insight into the history of the species. It also provides valuable direction to breeders in need to diversify their crop gene pools (28). Here we report a study revealing population structure and LD pattern of the U.S. core collection with good representation of the genetic diversity present in global rapeseed/canola germplasm accessions.

## Materials and methods

### Plant samples and phenotyping

A collection of 383 canola germplasm accessions originated in 24 countries comprising spring, winter, semi-winter, and rutabaga types, were collected from North Central Regional Plant Introduction Station (NCRPIS), Ames, Iowa, USA and North Dakota State University (NDSU) (S1 Table). The collection consists of 156 spring, 152 winter, 58 semi-winter, and 17 rutabaga types. Growth habit was determined by growing the accessions in a greenhouse for at least two seasons and in a field for two years at two locations. Field flowering time was recorded as number of days from seeding date to flowering date where first flowers opened on half of the plants belonging to a single accession. Spring type accessions flowered within 40-60 days after planting, the semi-winter types flowered from 70 to 110 days. On the other hand, winter types did not flower under field conditions. Vernalization treatment was conducted on greenhouse grown winter type accessions to induce flowering and also to confirm the winter habit type according to Rahman and McClean (2013) (6).

### Genotyping and sequencing

Young leaves were collected from 30 days old plants and flash-frozen in liquid nitrogen. Tubes were stored at -80° C until lyophilized. The lyophilized leaf tissue was ground in tubes with stainless beads using a plate shaker. DNA was extracted using Qiagen DNeasy Kit (Qiagen, CA, USA) from lyophilized tissue following the manufacturer’s protocol. DNA concentration was measured using a NanoDrop 2000/2000c Spectrophotometer (Thermofisher Scientific). The ApekI enzyme was used for GBS library preparation (29). Sequencing of the library was done at the University of Texas Southwestern Medical Center, Dallas, Texas, USA using Illumina HiSeq 2500 sequencer.

### SNP calling

TASSEL 5 GBSv2 pipeline (30) was used for SNP calling using a 120-base kmer length and minimum kmer count of ten. The reads were aligned to the canola reference genome (31) (available at: ftp.ncbi.nlm.nih.gov/genomes/all/GCF/000/686/985/GCF_000686985.2_Bra_napus_v2.0/) using Bowtie 2 (version 2.3.0) alignment tool (32). After passing all the required steps of TASSEL 5 GBSv2 pipeline, 497336 unfiltered SNPs were identified. Then VCFtools (33) was used to select bi-allelic SNPs considering the criteria: minor allele frequency (MAF) ≥ 0.05, missing values (max-missing) ≤50%, depth (minDP) ≥ 5 and physical distance (thin) ≤ 1000 bp. The SNPs that were located outside chromosomes (i.e. position unknown), were removed. As canola is a self-pollinating crop, the SNPs that were heterozygous in more than 25% of total genotypes, were also removed using TASSEL (34). All these filtering steps resulted in a total of 8,502 SNP markers.

### Data analysis

The core collection was divided into genetic groups using STRUCTURE v2.3.4 (35) software. The admixture model, a burnin period of 10000 and 50000 Monte Carlo Markov Chain (MCMC) iterations with 10 replications per K (K1-K10), were used as parameters for structure analysis. The optimal number of groups was determined based on DeltaK approach (36) which was performed by Structure Harvester (37). The individual Q matrix for the optimal K value was generated utilizing membership coefficient matrices of ten replicates from STRUCTURE analysis using CLUMPP (38). The results of structure analysis was visualized using the Structure Plot v2 software (39). Principal component analysis (PCA) was conducted by covariance standardized approach in TASSEL (34). An unrooted neighbor-joining (NJ) phylogenetic tree was constructed using MEGAX program with 1000 bootstraps (40). Resulting tree was displayed using FigTree V1.4.4 (41).

Analysis of molecular variance (AMOVA) was done to partition the genetic variance among the groups identified by STRUCTURE in Arlequin3.5. The average pairwise between-population *F_st_* values were also calculated using Arlequin3.5 (42). GenAlex v6.5 (43) was used to estimate percentage of polymorphic loci, number of effective alleles, Shannon’s information index, expected heterozygosity and unbiased expected heterozygosity of each marker and subpopulation. The SNP distribution plot was developed using R package CMplot (available at: https://github.com/YinLiLin/R-CMplot). The polymorphism information content (PIC) of markers was calculated using software Cervus (44). Tajima’s D value of each group was calculated using MEGAX software (40). The kinship (IBS) matrix was calculated using software Numericware i (45), kinship heatmap and histogram were developed using R package ComplexHeatmap (46). The level of relatedness (IBS coefficients) was correlated with Shannon’s information index (I) and diversity (H) in R v3.5.2 (47). Linkage disequilibrium (LD) pattern of whole collection and different subpopulations were analyzed using PopLDdecay (48). The mean linked LD was calculated by dividing total *r^2^* value with total number of corresponding loci pair. In this case, *r^2^* < 0.2 was considered only. Same procedure was followed to calculate mean unlinked LD where *r^2^* ≥ 0.2 was considered. Haplotype block analysis was done using PLINK (49) with a window size of 5 Mb. Confidence interval (CI) method (50) was used to identify haplotype blocks with high LD. Haplotype blocks (>19 Kb), observed in one subpopulation but not in the other, were considered to be subpopulation-specific block. Haplotype blocks (>19 Kb) shared by more than one subpopulation, were considered to be common to corresponding subpopulations.

## Results

### SNP profile

The selected 8,502 SNPs were distributed across 19 chromosomes with an average marker density of 1 per 99.5 kb. Chromosome A3 and A4 contained highest (685 SNPs, 8.06%) and lowest (236 SNPs, 2.78%) number of SNPs, respectively. The SNP density was highest on chromosome A7 (71.1 kb) and was lowest on chromosome C9 (134.5 kb) (Table 1, Fig 1). The occurrence of transition SNPs (4,956 SNPs) was more than that of transversions (3,546 SNPs) with a ratio of 1.40. The ratio of transitions to transversions SNPs was higher in A genome (1.41) than that of in C genome (1.38). In both genome, G/C transversions were lowest (4.33% and 4.29%), but A/G and C/T transitions occurred in almost similar frequencies (Table 2). The inbreeding coefficient within individuals (*F_st_*), inbreeding coefficient within subpopulations (*F_st_*), observed heterozygosity (*Ho*) and fixation index (*F*) of all the markers ranged from -0.45 to 1.00, 0 to 0.73, 0 to 0.57 and 0.40 to 1.00, respectively. The Shannon’s information index (*I*) of all markers ranged from 0.10 to 0.69 with a mean value of 0.37. The expected heterozygosity (*He*) ranged from 0.05 to 0.50 with a mean value of 0.27. The polymorphic information content (*PIC*) ranged from 0.05 to 0.37 with a mean value of 0.22 (S2 Table). Subpopulation-wise marker diversity parameters are presented in supplementary S3 Table.

**Fig 1.**
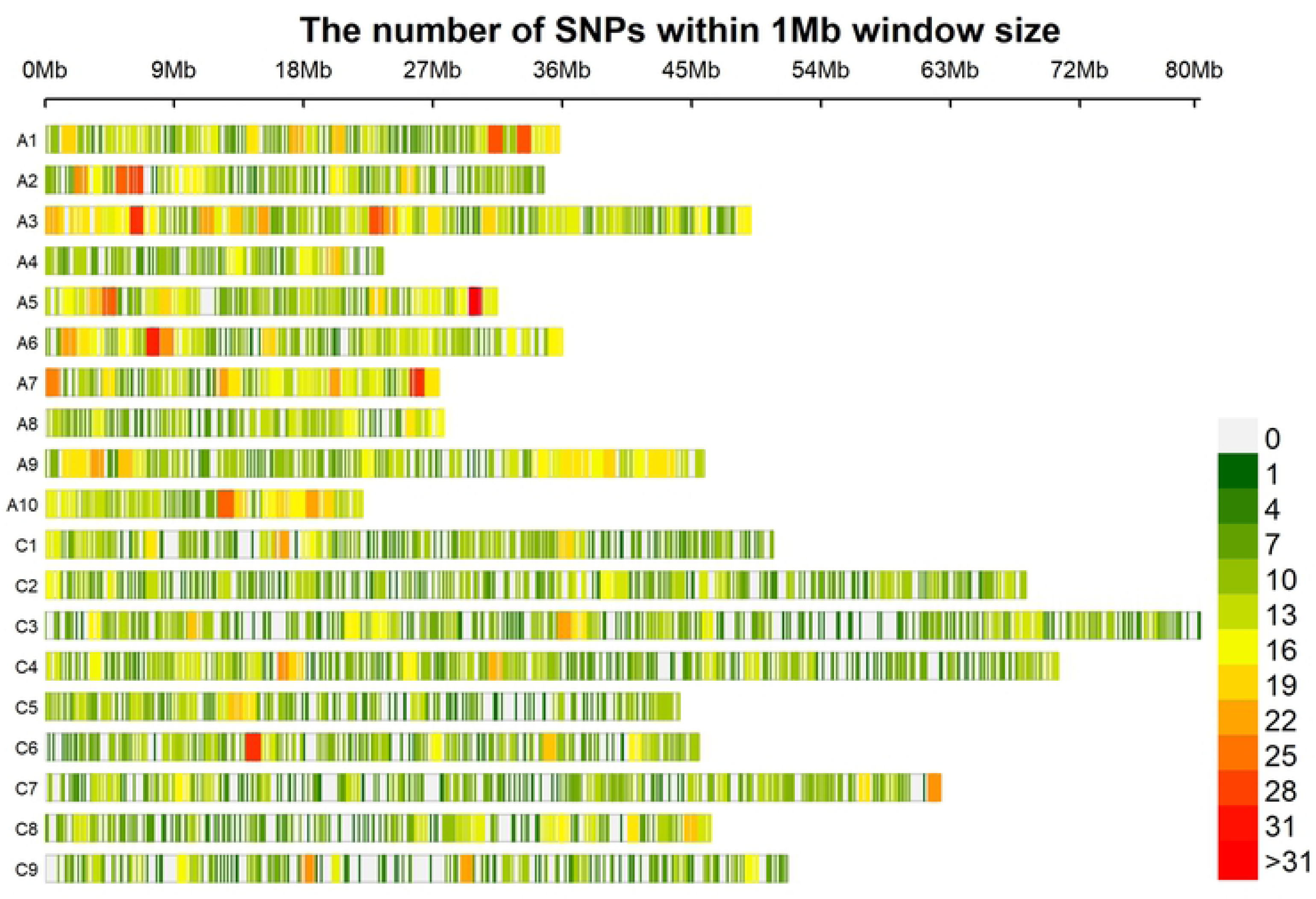
Chromosome-wise SNP density map. Frequency of SNPs varies according to color gradient.

**Table 1.** Chromosomewise distribution of SNP markers.

| Chromosome | No. of SNPs | % SNPs | Start position | End position | Length (Mb) | Density (Kb) |
| --- | --- | --- | --- | --- | --- | --- |
| A1 | 440 | 5.18 | 149163 | 35806075 | 35.7 | 81.0 |
| A2 | 392 | 4.61 | 13430 | 34692905 | 34.7 | 88.5 |
| A3 | 685 | 8.06 | 2769 | 49103583 | 49.1 | 71.7 |
| A4 | 236 | 2.78 | 32805 | 23517671 | 23.5 | 99.5 |
| A5 | 413 | 4.86 | 18668 | 31435105 | 31.4 | 76.1 |
| A6 | 448 | 5.27 | 120409 | 36005103 | 35.9 | 80.1 |
| A7 | 384 | 4.52 | 85869 | 27388322 | 27.3 | 71.1 |
| A8 | 281 | 3.31 | 231427 | 27734410 | 27.5 | 97.9 |
| A9 | 541 | 6.36 | 81404 | 45841268 | 45.8 | 84.6 |
| A10 | 305 | 3.59 | 133853 | 22085737 | 22.0 | 72.0 |
| C1 | 445 | 5.23 | 86671 | 50660872 | 50.6 | 113.7 |
| C2 | 589 | 6.93 | 92431 | 68260222 | 68.2 | 115.7 |
| C3 | 651 | 7.66 | 3839 | 80365889 | 80.36 | 123.4 |
| C4 | 634 | 7.46 | 138930 | 70507417 | 70.4 | 111.0 |
| C5 | 366 | 4.30 | 26760 | 44124497 | 44.1 | 120.5 |
| C6 | 414 | 4.87 | 275190 | 45479327 | 45.2 | 109.2 |
| C7 | 518 | 6.09 | 271113 | 62304827 | 62.0 | 119.8 |
| C8 | 383 | 4.50 | 57934 | 46317429 | 46.3 | 120.8 |
| C9 | 377 | 4.43 | 920885 | 51627086 | 50.7 | 134.5 |
| Mean | 447.47 |  |  |  |  | 99.5 |

**Table 2.** Transition and transversion SNPs across the genome.

| Genome | SNP type | Model | No. of sites | Frequencies (%) | Total (percentage) |
| --- | --- | --- | --- | --- | --- |
| A | Transitions | A/G | 1195 | 14.06 | 2416 (28.3%) |
|  |  | C/T | 1221 | 14.36 |  |
|  | Transversions | A/T | 457 | 5.38 | 1709 (20.1%) |
|  |  | A/C | 424 | 4.99 |  |
|  |  | G/T | 460 | 5.41 |  |
|  |  | G/C | 368 | 4.33 |  |
| C | Transitions | A/G | 1273 | 14.97 | 2540 (29.9%) |
|  |  | C/T | 1267 | 14.90 |  |
|  | Transversions | A/T | 496 | 5.83 | 1837(21.6%) |
|  |  | A/C | 482 | 5.67 |  |
|  |  | G/T | 494 | 5.81 |  |
|  |  | G/C | 365 | 4.29 |  |

### Population structure

The whole collection was divided into six subpopulations based on structure analysis using the Delta K approach (Fig 2A). Genotypes of different types and origins were well clustered. The winter, semi-winter, rutabaga type genotypes were grouped under subpopulation-1 (P1), subpopulation-2 (P2), and subpopulation-5 (P5), respectively. The spring type genotypes were grouped under two subpopulations: subpopulation-3 (P3) and subppulation-4 (P4) whereas subpopulation-6 (P6) composed of only nine genotypes of different types (Fig 2B). Though each subpopulation consisted of genotypes of different origin, P1, P2 and P4 were dominated by European, Asian and American (NDSU breeding lines) genotypes, respectively, whereas P3, P5 and P6 were composed of genotypes of mixed origin. We performed principal component analysis (PCA) to show the genetic similarity among subpopulations and genotypes. The first two axes explained 21% of the total observed variation (S4 Table). The PCA revealed that subpopulation P1, P2, P3, P4 and P5 were well clustered and separated from each other, but genotypes of P6 were scattered within other subpopulations (Fig 3). In addition to that, we also constructed phylogenetic tree based on neighbor joining (NJ) criteria (Fig 4). The output of neighbor-joining (NJ) tree analysis was in line with that of PCA. Based on the PCA and NJ output, we merged the P6 genotypes into P1, P2, P3 and P4 and considered five subpopulations for further analysis and discussions. Based on individual Q matrix, the proportion of pure (non-hybrid) and admixed (containing markers assigned to more than one subpopulation) genotypes in each subpopulation was calculated. The proportion of pure accessions in subpopulations ranged from 41% to 80% at a 0.7 cutoff value and 10% to 35% at 0.9 cutoff value (Table 3).

**Fig 2.**
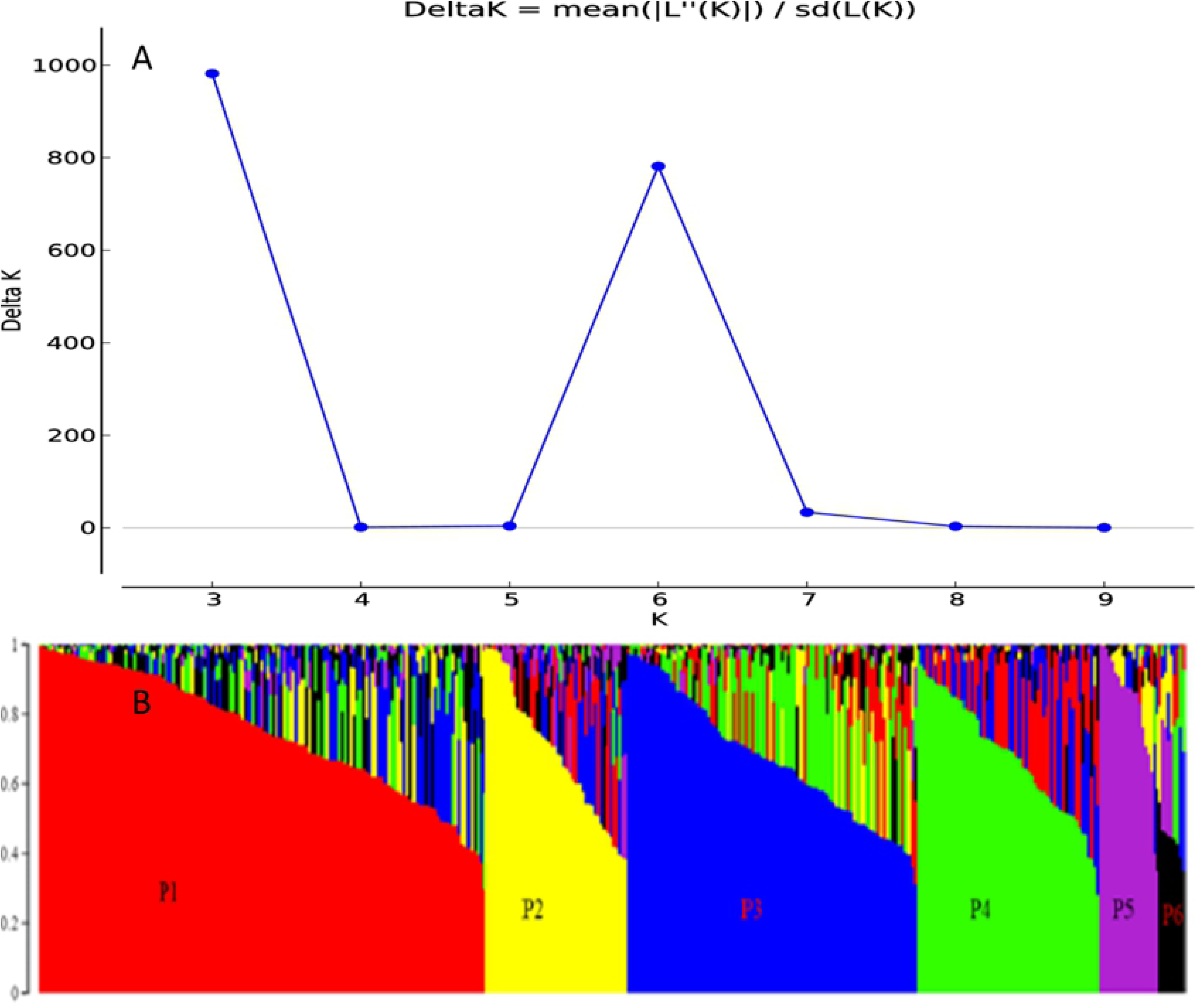
Bayesian clustering of whole collection using 8,502 SNP markers in STRUCTURE v.2.3.4. (A) Graphical representation of optimal number of clusters (K) determined by Evanno’s method, where highest Delta K indicate the number of subpopulations. (B) Estimated population structure (P1 to P6) of 383 canola genotypes on K=6 according to Delta K.

**Fig 3.**
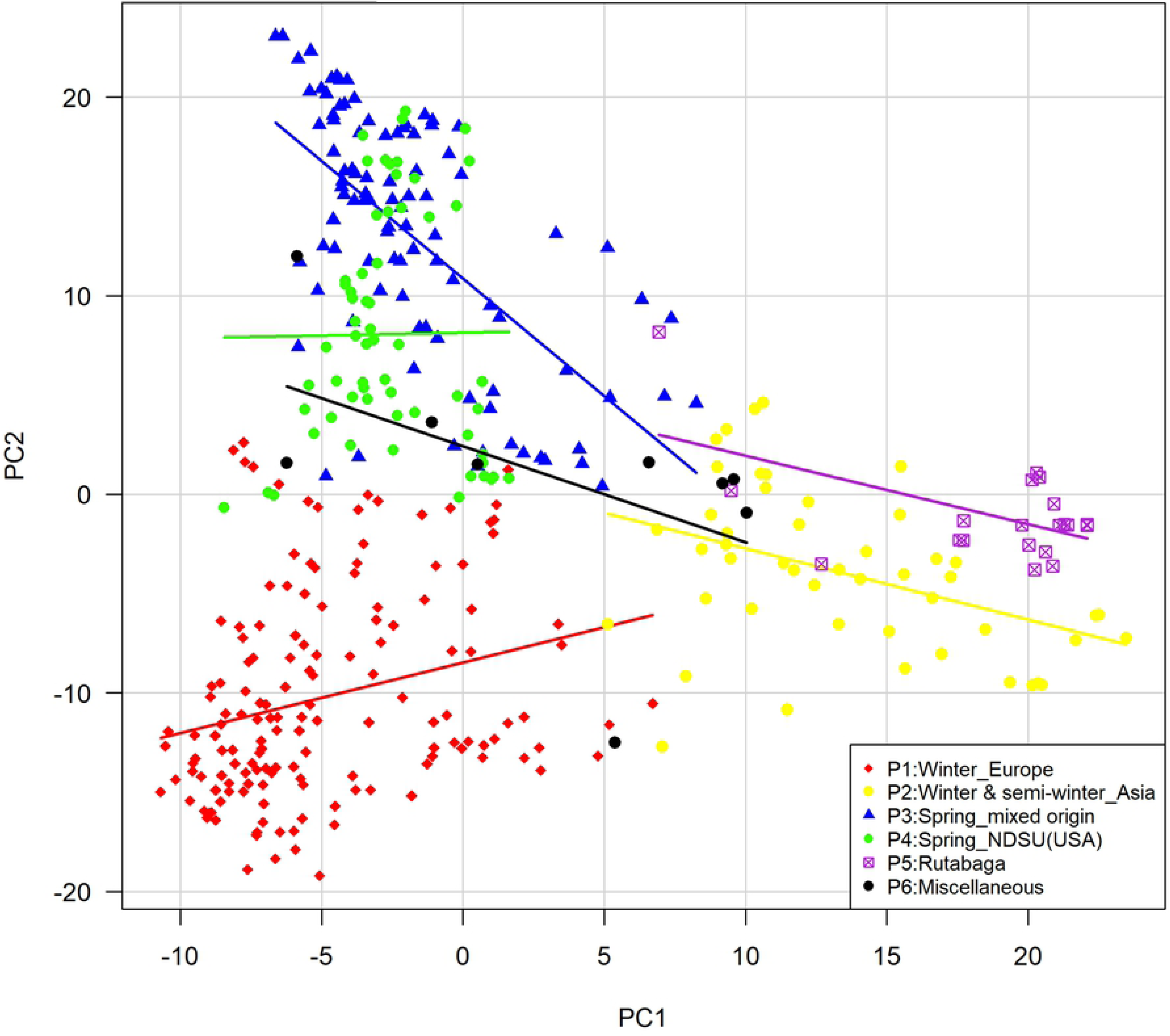
Principal component analysis of SNP diversity based on genetic distance. Colors represent different subpopulations identified at K = 6 in figure 2. Genotypes belong to P6 (black dots) are scattered within different subpopulations.

**Fig 4.**
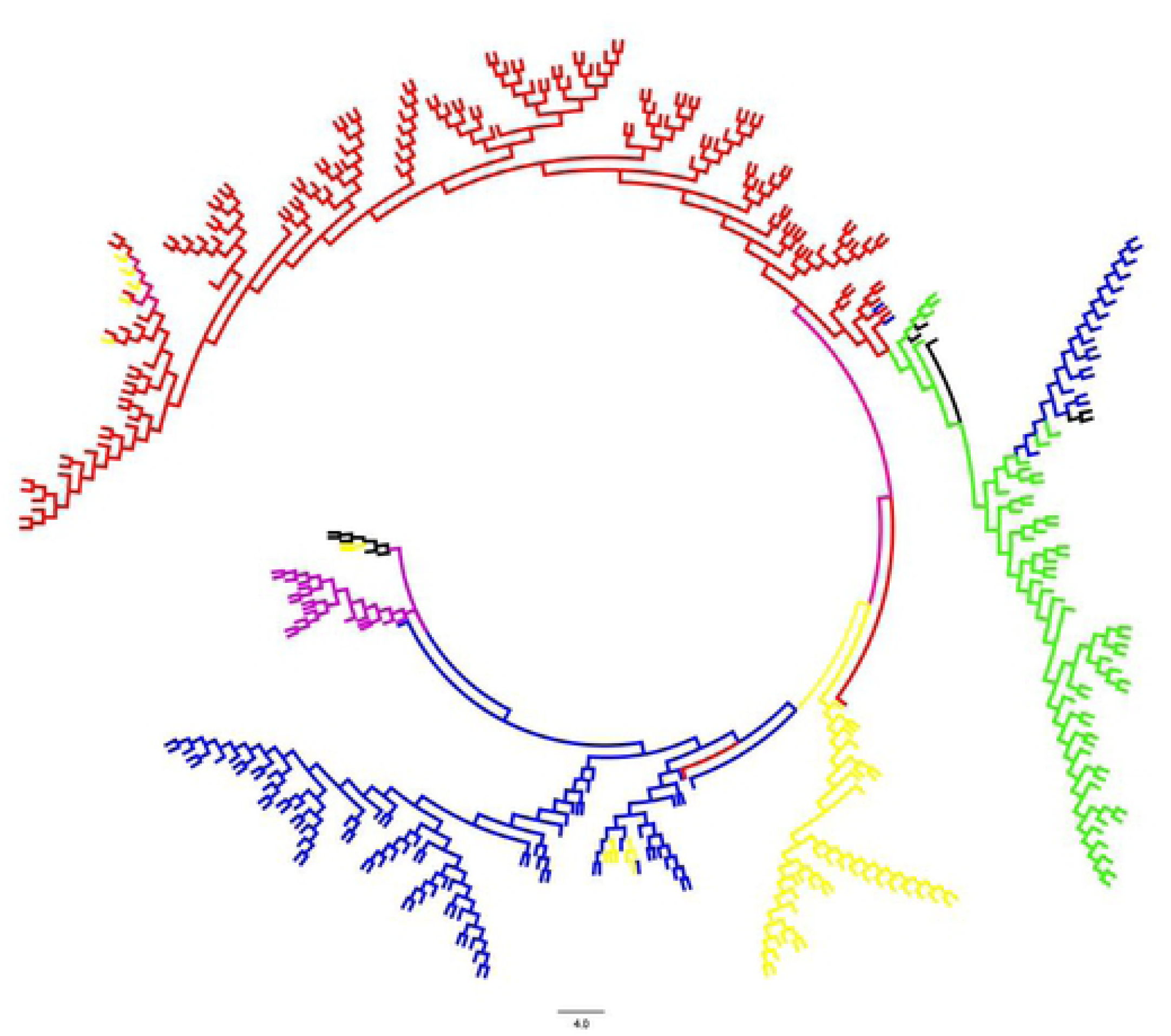
Phylogenetic tree based on neighbor-joining (NJ) using information from 8,502 SNP markers. Each branch is color-coded according to membership into the K= 6 subpopulations identified by structure in figure 2. Subpopulations bear same color as in figure 3.

**Table 3.**
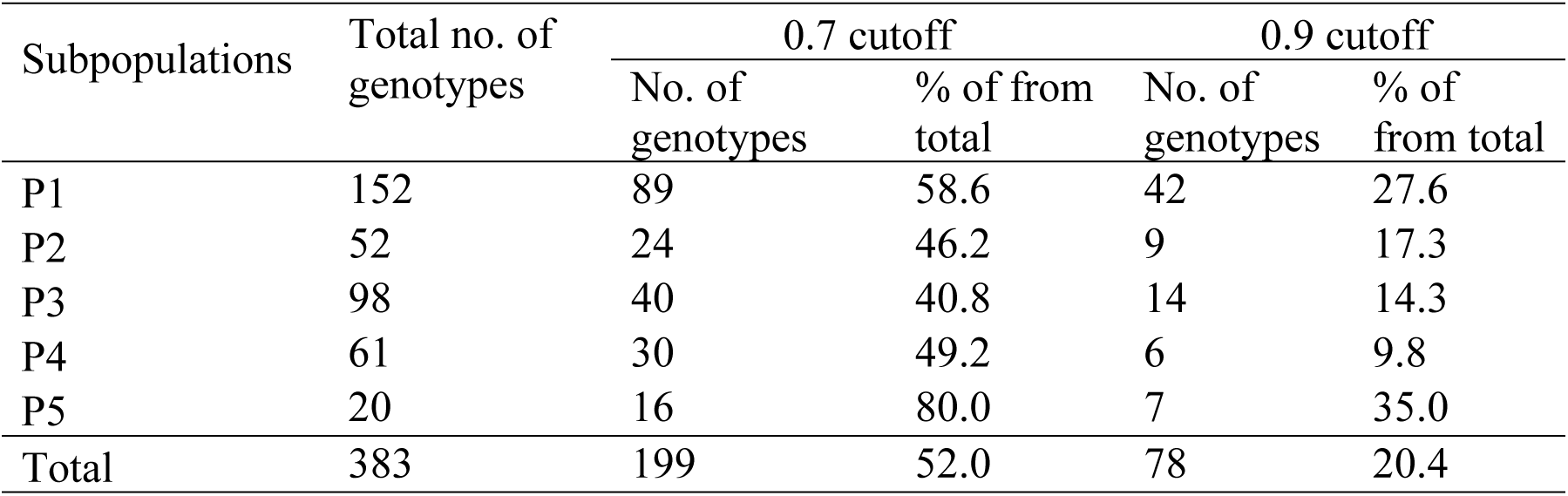
Number of pure and admixed individuals per subpopulation.

| Subpopulations | Total no. of genotypes | 0.7 cutoff |  | 0.9 cutoff |  |
| --- | --- | --- | --- | --- | --- |
|  |  | No. of genotypes | % of from total | No. of genotypes | % of from total |
| P1 | 152 | 89 | 58.6 | 42 | 27.6 |
| P2 | 52 | 24 | 46.2 | 9 | 17.3 |
| P3 | 98 | 40 | 40.8 | 14 | 14.3 |
| P4 | 61 | 30 | 49.2 | 6 | 9.8 |
| P5 | 20 | 16 | 80.0 | 7 | 35.0 |
| Total | 383 | 199 | 52.0 | 78 | 20.4 |

### Population diversity

In all subpopulations, the percentage of polymorphic loci was greater than 75%. It was highest in P1 (99%) and lowest in P5 (75%). The diversity (*H*) of the five subpopulations ranged from 0.19 (P4 and P5) to 0.25 (P2) with an average of 0.22. The Shannon’s information index (*I*) ranged from 0.31 (P4 and P5) to 0.40 (P2) with an average of 0.34. The Tajima’s D value ranged from -0.70 (P4) to 0.53 (P1) with an average of 0.13 (Table 4).

**Table 4.** Subpopulation-wise diversity parameters.

| Subpopulations | Polymorphic loci (%) | $N_a$<sup>a</sup> | $N_e$<sup>b</sup> | $I$<sup>c</sup> | $H$<sup>d</sup> | $U_h$<sup>e</sup> | Tajima's D* |
| --- | --- | --- | --- | --- | --- | --- | --- |
| P1 | 99.12 | 1.99 | 1.32 | 0.35 | 0.21 | 0.21 | 0.53 |
| P2 | 94.32 | 1.94 | 1.40 | 0.40 | 0.25 | 0.25 | 0.30 |
| P3 | 96.98 | 1.97 | 1.35 | 0.36 | 0.22 | 0.23 | 0.30 |
| P4 | 80.67 | 1.81 | 1.30 | 0.31 | 0.19 | 0.19 | -0.70 |
| P5 | 75.25 | 1.75 | 1.31 | 0.31 | 0.19 | 0.20 | 0.23 |
| Mean | 89.27 | 1.89 | 1.34 | 0.34 | 0.22 | 0.22 | 0.13 |

<sup>a</sup> No. of different alleles, <sup>b</sup> No. of effective alleles, <sup>c</sup> Shannon's information index
<sup>d</sup> Diversity, <sup>e</sup> Unbiased diversity. SE (standard error) was zero in all cases. Indices calculated using 8191 SNPs with GenAlex v. 6.5. \* was calculated with 1000 permutations.

### Population genetic differentiation

The analysis of molecular variance (AMOVA) revealed that variance among subpopulations covered 24% of total variation whereas the remaining 76% of total variation accounted for variance among individuals within subpopulations (Table 5) with a *F_st_* and *Nm* value of 0.24 and 1.28, respectively.

**Table 5.** Summary of AMOVA.

| Sources of variation | d.f. | Sum of squares | Variance components | % of variation | $F_{st}$ | $Nm$ |
| --- | --- | --- | --- | --- | --- | --- |
| Among subpopulations | 4 | 130814.6 | 228.5*** | 23.5 | 0.24 | 1.28 |
| Within subpopulations | 761 | 565699.6 | 743.4 | 76.5 |  |  |
| Total | 765 | 696514.1 | 971.9 |  |  |  |

\*\*\* indicates $p < 0.001$ for 1023 permutations

All pairwise *F_st_* comparisons between subpopulations were significant (*p* < 0.01). All combinations showed a great degree of divergence (*F_st_* > 0.20) (Wright 1943), except combinations P3 and P4 (0.11), P1 and P2 (0.19). The pairwise *F_st_* > 0.30 was observed between P1 and P5, P3 and P5, P4 and P5 (Table 6).

**Table 6.** Genetic differentiation among subpopulations.

| Subpopulation pairwise $F_{st}$ | | | | | |
| --- | --- | --- | --- | --- | --- |
|  | P1 | P2 | P3 | P4 | P5 |
| P1 | 0 |  |  |  |  |
| P2 | 0.19** | 0 |  |  |  |
| P3 | 0.25** | 0.24** | 0 |  |  |
| P4 | 0.21** | 0.24** | 0.11** | 0 |  |
| P5 | 0.34** | 0.24** | 0.34** | 0.39** | 0 |

Diagonal values are pairwise $F_{st}$ comparisons, performing 1000 permutations using Arlequin v.
3.5. \*\*indicates $p < 0.01$

We performed kinship (IBS) analysis to facilitate the individual genotype selection for desirable cross combinations (Fig 5, S5 Table). In whole collection, the IBS coefficients ranged from 1.21 to 1.94. The average coancestry between any two canola genotypes was 1.47. The P2 contained almost 50% of total genotypic pairs having IBS coefficients less than 1.50 whereas this portion was comparatively lower in other subpopulations (Table 7, S1 Fig).

**Fig 5.**
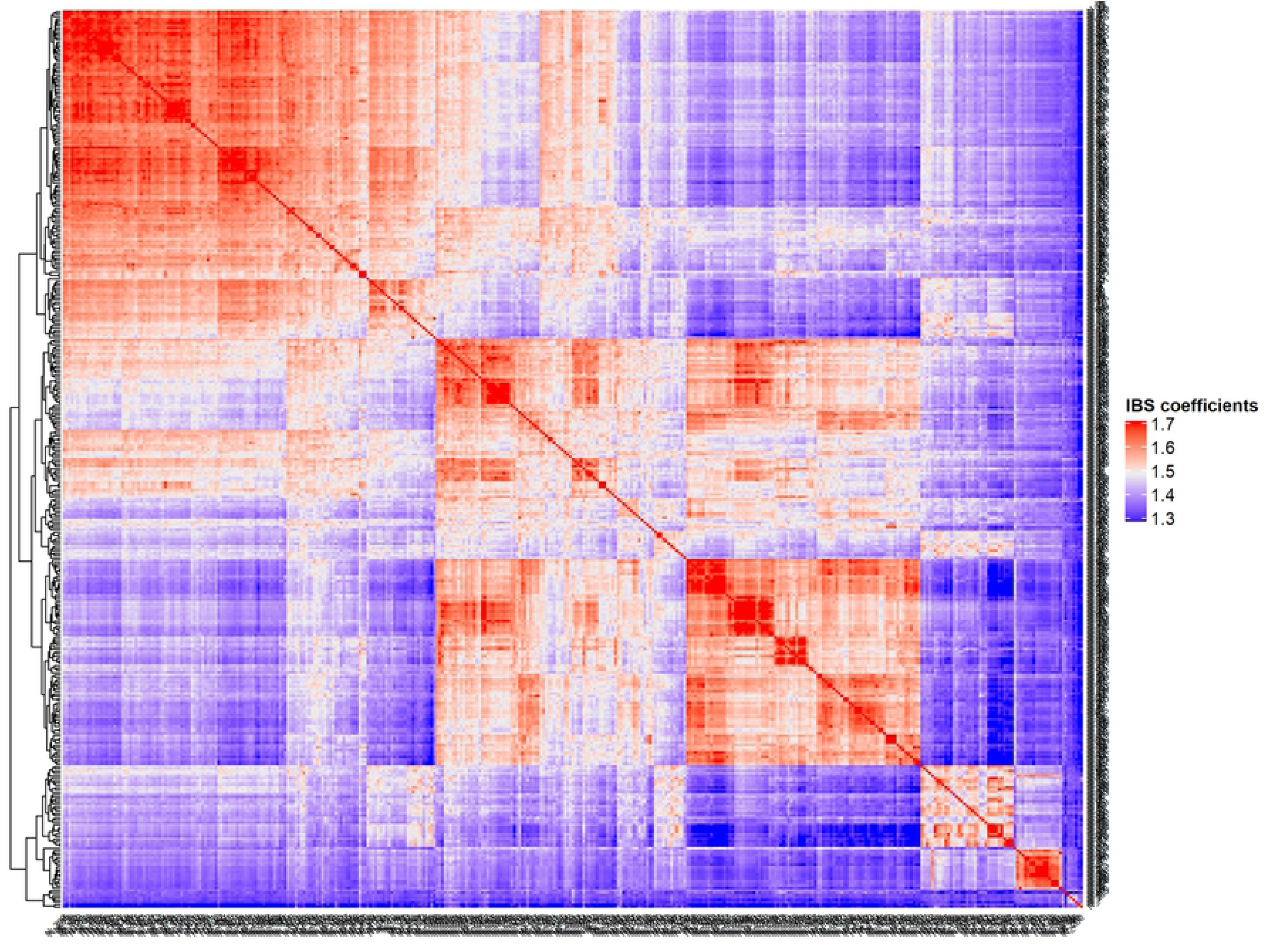
Heatmap of kinship matrix of entire collection.

**Table 7.** Summary of subpopulation-wise kinship (IBS) matrix.

| Subpopulations | Whole collection | P1 | P2 | P3 | P4 | P5 |
| --- | --- | --- | --- | --- | --- | --- |
| IBS coefficients range | 1.21- 1.94 | 1.40- 1.94 | 1.27- 1.93 | 1.29- 1.93 | 1.46-1.94 | 1.35-1.92 |
| Mean of IBS coefficients | 1.47 | 1.58 | 1.49 | 1.55 | 1.62 | 1.60 |
| Pairs having $\leq 1.50$ IBS coefficients (%) | 63.9 | 9.6 | 50.7 | 21.7 | 1.1 | 18.0 |
| Pairs having $> 1.50$ IBS coefficients (%) | 36.1 | 90.4 | 49.3 | 78.3 | 98.9 | 82.0 |

We also performed correlation analysis between mean pairwise relatedness (IBS coefficients) among individuals within subpopulation and Shannon’s information index (*I*), diversity (*H*). The *I* and *H* were significantly and negatively correlated with relatedness (*r*= -0.97, -0.98, and *p*<0.01), respectively.

### Linkage disequilibrium pattern

The linkage disequilibrium (LD) pattern was investigated across the entire collection, each subpopulation, genome, and chromosome-wise. LD = *r^2^* values decreased with the increase of distances. In all cases, mean LD was high (*r^2^* > 0.22) at short distance bin (0-2 kb) and declined with increasing bin distance (S6 Table). In the entire collection considering both A and C genome, the mean linked LD, mean unlinked LD and loci pair under linked LD was 0.44, 0.02 and 1.81%, respectively. The mean linked LD was similar in P3 and P4 (*r^2^* = 0.45), lower in P2 (*r ^2^*= 0.41) and higher in P1 (*r ^2^*= 0.48). The loci pair in linked LD was higher in P5 (8.76%) and lowest in P1 (1.52%). The mean linked LD, mean LD and loci pair under linked LD was always higher in all cases in case of C genome than that of A genome (Table 8). We also calculated the LD decay rate. In the whole collection, LD decayed to its half maximum within < 45 kb distance for whole genome, < 21 kb for A genome, and < 93 kb for C genome. In all subpopulations, the distance for LD decay to its half maximum was always higher for C genome than A genome. Each chromosome showed differential rate of LD decay (S2 and S3 Fig). LD persisted the longest in chromosome C1 (348 kb) and C2 (244 kb). The decay distance was shortest in chromosome A5 (13 kb) and A1 (16 kb) (S6 Table). LD decayed to its half –maximum within < 29 kb for P1, <45 kb for P2, P3, <101 kb for P4, and <120 kb for P5. In all subpopulations LD persisted also longest in all chromosomes of C genome than that of A genome (Fig 6, S7 Table).

**Fig 6.**
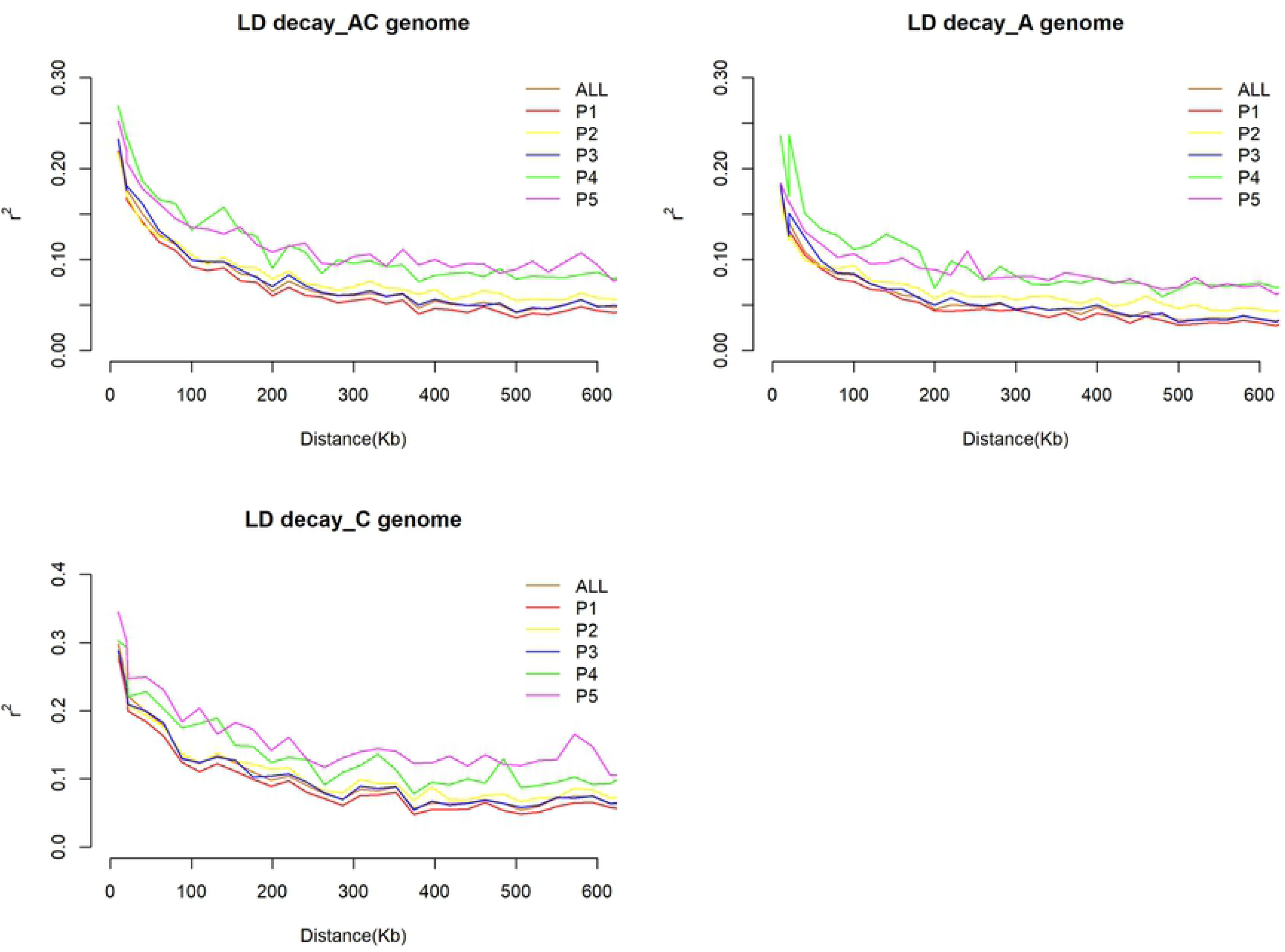
Linkage disequilibrium (LD) differences and decay pattern among subpopulations.

**Table 8.**
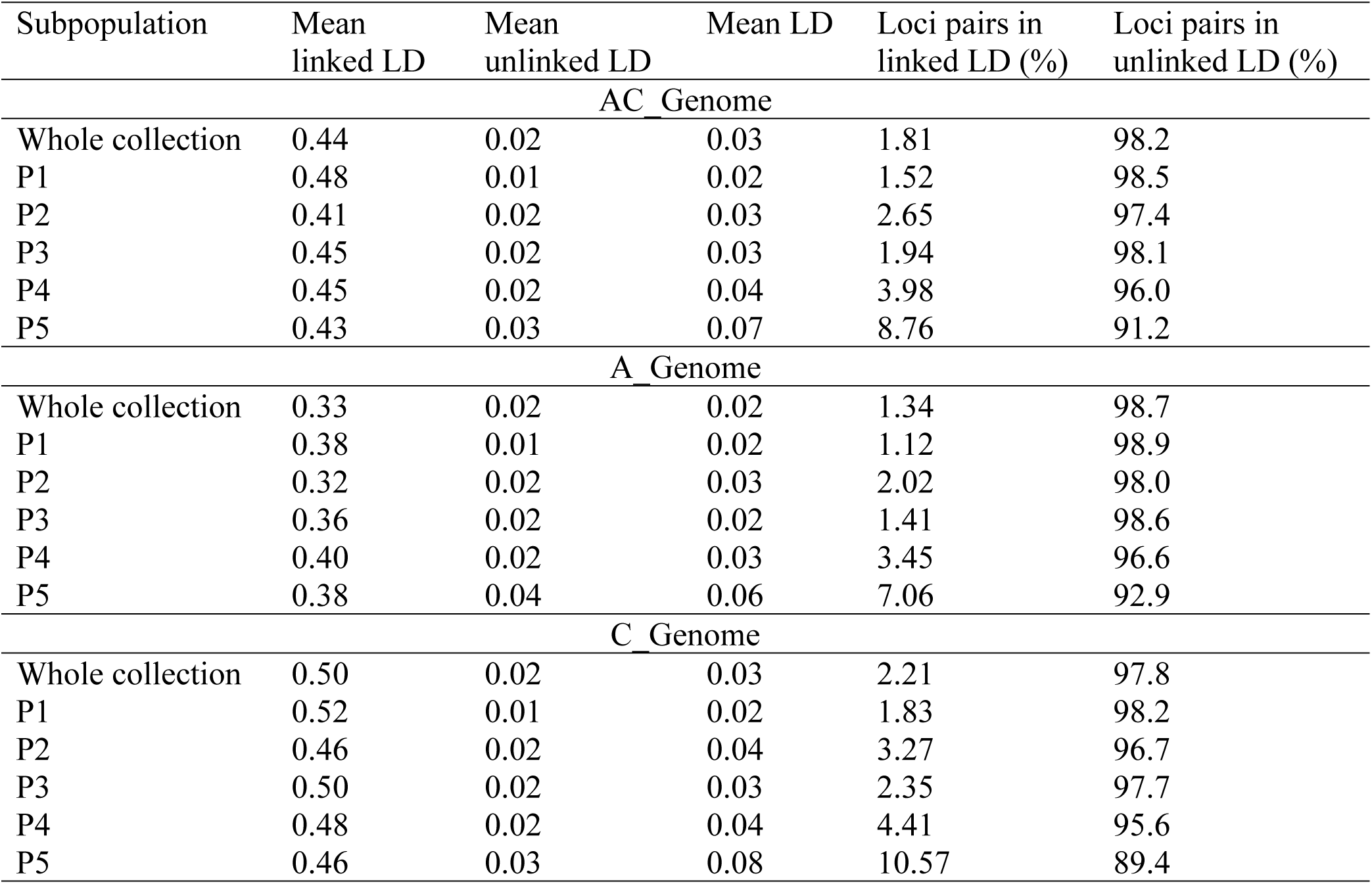
Linkage disequilibrium in the studied collection.

We also performed haplotype block analysis to investigate LD variation patterns across whole genome. A total 200 blocks covering 18 Mb out of the 976 Mb anchored *B. napus* reference genome (31), were identified. A and C genome contained 67 and 133 haplotype blocks, respectively. The total length of A and C genome specific haplotype blocks were 1.8 Mb and 16 Mb, respectively. The total length of haplotype blocks varied greatly from chromosome to chromosome. Total haplotype block length varies from 24 Kb on A1 to 901 Kb on A9 in A genome and in C genome it varies between 40 Kb on C9 to 3,610 Kb on C2 (Table 9). The subpopulation-specific and common haplotype blocks were shown in supplementary S8 Table.

**Table 9.**
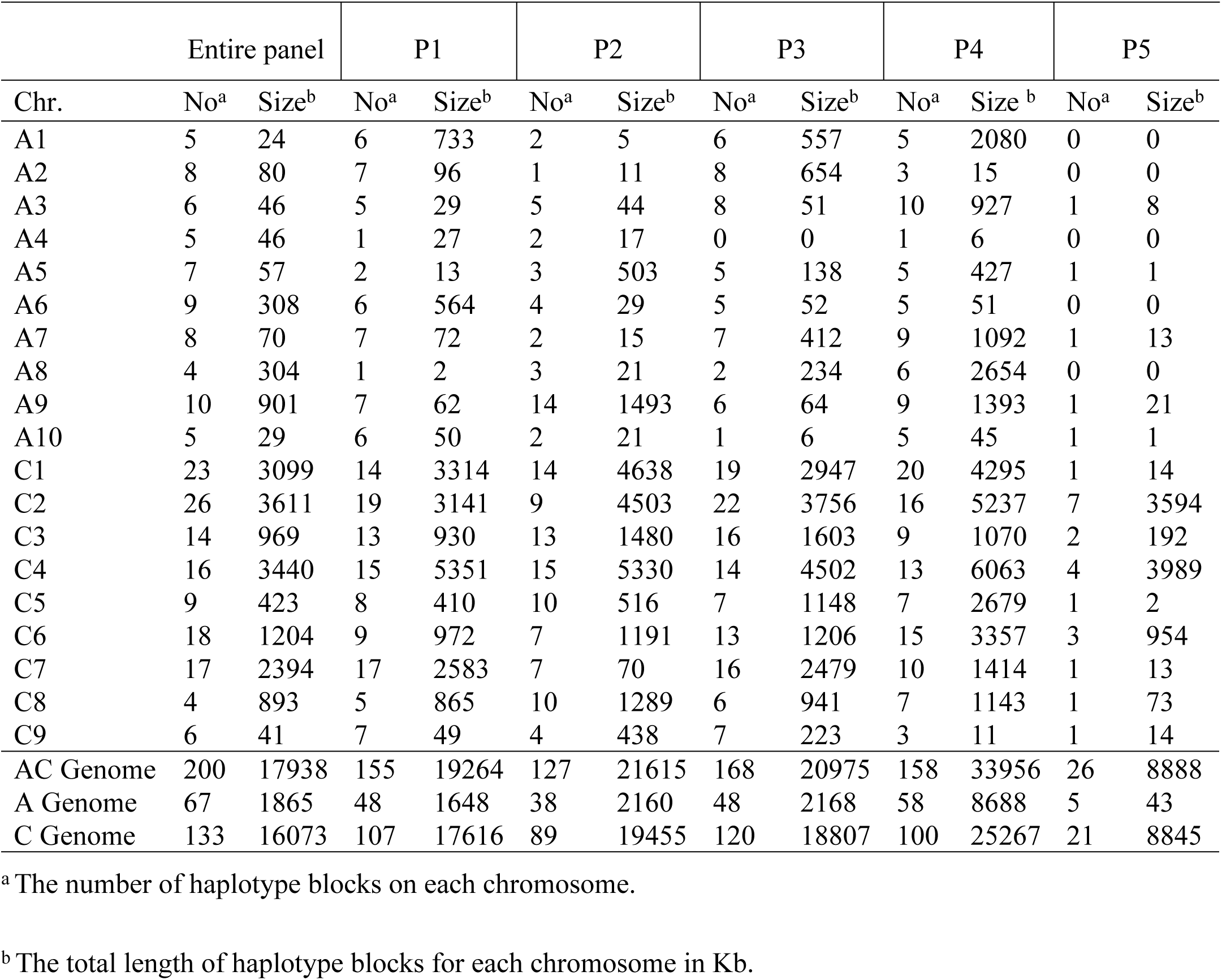
Subpopulation-wise number and length of haplotype blocks (HBs) along chromosomes.

## Discussion

Genotyping-by-sequencing (29) is one approach to obtain high frequency SNPs. The strategy has been used for population genetic studies, association mapping, and proven to be a powerful tool to dissect multiple genes/QTL in many plant species (51–53). We obtained 497,336 unfiltered SNPs markers of which 8,502 high quality SNP marker were used for genetic diversity analysis of 383 genotypes. Delourme et al. (2013) (24) conducted genetic diversity analysis in *B. napus* using 7,367 SNP markers of 374 genotypes. However, different marker technologies such as Single Sequence Repeat (SSR), Sequence Related Amplified Polymorphism (SRAP) markers have been used by other researchers for genetic diversity analysis in *B. napus*. Chen et al. (2020) (54) used 30 SSR markers, Wu et al. (2014) (55) utilized 45 SSR markers, Ahmad et al. (2014) (56) used 20 SRAP markers for genetic diversity and population structure analysis of *B. napus*. Earlier, our group conducted a genetic diversity study of flax using 373 germplasm accessions with 6,200 SNP markers.

The SNP markers were distributed throughout 19 chromosomes of *B. napus* and the marker density was 1 per 99.5 Kbp. This is comparable density to earlier study conducted by Delourme et al. (2013) (24). Therefore, this marker density provides a sufficient resolution to accurately estimate genome-wide diversity as well as the extent of LD within the genome. The high marker density will also help association mapping studies precisely identify a causal locus/loci or very closely linked loci that can be further used either in MAS or to pinpoint the causative locus (57).

The core collection utilized in this study represents mostly adapted lines from various breeding programs rather than wild accessions commonly used in diversity studies. Therefore, sources of variation, markers of interest identified in the collection can be directly used in breeding programs.

We have identified higher frequency of transition SNPs over transversion SNPs that is an agreement with Bus et al. (2012) (58), Clarke et al. (2013) (59), and Huang et al. (2013) (60) in *B. napus*. Higher number of transition SNPs over transversion is also reported in other crop species such as *Hevea brasiliensis* (61), *Camellia sinensis* (62), *Camelina sativa* (63), and *Linum usitatissimum* (64). The high number of transition SNPs indicating that this mutation is more tolerable to natural selection (65).

We have calculated the polymorphic information content (PIC) and expected heterozygosity (*He*) for each marker. The PIC indicates the usefulness of any marker for linkage analysis, and *He* determines the diversity of haploid markers (66). In our research, the PIC value is ranged from 0.05 to 0.35 indicating that the markers are moderate or low informative. The similar lower PIC value (0.1 to 0.35) was reported by Delourme et al. (2013) (24) in *B. napus*. The lower PIC value was also reported in winter wheat (67), maize (68), flax (64, 69) and rice (70). The lower PIC value is a result of bi-allelic nature of SNP markers and probable low mutation rate (71). In our study, the *He* value of each marker was always greater than corresponding PIC value indicating an average lower allele frequency in our population (66).

### Population structure and diversity

Multiple population-based analyses, including population structure, PCA and NJ tree analysis split the core collection into five distinct clusters. Previously, three clusters have been reported based on winter, spring and semi-winter growth habit types of *B. napus* (20, 24–26,72–74). Our analysis showed a clear trend of the five clusters belonging to winter type (P1), semi-winter type (P2), spring type of mixed origin (P3), spring type developed at NDSU (P4), and rutabaga type (P5) growth habits. Our finding showed strong agreement with other studies to separate winter, semi-winter and spring types. Moreover, we could separate the rutabaga types and divided the spring type into two groups: American origin and the rest of other countries origin. This additional cluster added new information to select germplasm from different subpopulation for increasing genetic diversity in canola breeding program.

We have identified a moderate diversity (average *H* = 0.22) within the subpopulations. *B. napus* is capable of self-pollination, and little cross pollination may be occurred by insect. Being a mostly self-pollinated crop a low to moderate subpopulation diversity in *B. napus* is expected. Low to moderate diversity was also found in previous studies (75–77). Along with the reproduction system, one needs to look at evolution and domestication history for explaining low to moderate levels of diversity in *B. napus*. This allopolyploid species originated at Mediterranean coast as a result of a natural cross between *B. rapa* and *B. oleracea* which occurred approximately 0.12 -1.37 million years ago (78, 79). The domestication of *B. napus* occurred very recently, around 400 years ago with the first rapeseed being most likely a semi-winter type due to the mild climate in the region (72, 80). Later on, European growers developed the winter and spring type Brassicas through selection for cold hardiness or early flowering to expand its cultivation in further North in the last century (81). Therefore, the low to moderate diversity in winter and spring *B. napus* can be mostly explained by a recent history of the species, followed by infrequent exchange of genetic material with other Brassicas (24), as well as by the traditional breeding practices selecting for only few phenotypes. In our study, the more diversity in semi-winter type (P2, *H*= 0.25) than winter (P1, *H*= 0.21) and spring (P4, *H*= 0.19) type is supported by its domestication history. We have seen a homogeneity of the diversity indices of different subpopulations indicating that the species is stable enough to avoid the natural loss of genetic variability by genetic drift (82). The *Nm* value was greater than one, which indicates that there was enough gene flow among semi-winter, winter and spring types. These findings also support the evolution of winter and spring types from semi-winter type. In this research, Tajima’s D value was calculated to identify different subpopulations with availability or scarcity of rare alleles (83). P4 (spring type originated at NDSU, USA) showed a negative Tajima’s D value indicating presence of more rare and unique alleles and recent expansion of this subpopulation compared to the other subpopulations (84). Recently, the NDSU canola breeding program developed the P4 advanced breeding lines through crossing different genetic resources including winter, spring and semi winter types and subsequent selection. Therefore, the P4 displayed an abundance of rare alleles. The subpopulation P1, P2, P3, and P5 showed positive Tajima’s D value indicating an excess of intermediate frequency alleles, which may be caused by balancing selection, population bottleneck, or population subdivision. Previously, negative Tajima’s D values were found in spring and winter type B. napus accessions (74). The negative correlation between diversity indices (*H* and *I*) and relatedness (average IBS coefficients) indicates that inbreeding and genetic drift play a significant role in reducing genetic variability in the studied population which results in increased differentiation among sub-populations. Similar phenomenon was also found in flax (64) and *Arapaima gigas* species (85).

Parent selection for crossing from diverged population will allow us to create a new population with increased genetic diversity and transgressive segregation, and eventually will increase genetic gain. Pairwise *F_st_* statistic, a parameter describing population structure differentiation and the degree of divergence among populations (86), was estimated among five subpopulations. In the present study all pairwise *F_st_* values comprising both low and high values, were statistically significant. Similar type results were also found in other studies (74, 87–89). Lower pairwise *F_st_* (0.11) was identified between spring type originated in USA (P4) and spring type originated in other countries (P3). This is reasonably justified as both subpopulations comprise of spring type genotypes and germplasm exchanged occurred between USA and other countries. It also indicating that we will not get higher genetic diversity in population if we use only spring types in the crossing program. But this combination is good for accumulating specific elite trait if the targeted trait is found in members of one and missing from the members of another group. We found spring type (P3 and P4) genotypes are greatly divergent (*F_st_* > 0.20) from winter and semi-winter type (P1 and P2) genotypes. Utilization of genotypes from these group in crossing program will broaden the genetic base of developed population results in high heterosis. This potentiality has already been proved as hybrids between the Chinese semi-winter and European (including Canada) spring type exhibited high heterosis for seed yield (90). The P5 (rutabaga type) showed the higher *F_st_* with other subpopulations such as the highest *F_st_* was observed between P5 and P4 (NDSU spring type) followed by P3 (other spring type), P1 (winter type) and P2 (semi-winter type). This outcome clearly shows that rutabaga is genetically distinct from spring and winter type canola which is confirmed by previous studies (72, 73, 91). This distinctness of rutabaga can be exploited through heterosis breeding. Several previous studies have already showed rutabaga as a potential gene pool for the improvement of spring canola (92, 93). NDSU canola breeding program also utilized winter and rutabaga types in the breeding program for increasing genetic diversity and for improvement of spring canola.

Crossing among genotypes within subpopulations is also useful as variation among individuals within subpopulations is greater than that among subpopulations, which was revealed by AMOVA. This finding is also in agreement with the previous findings (54, 77, 94, 95). In this case subpopulations P2, P3 and P1 showing high diversity (*H* > 0.20) could be utilized for cultivar development. Structure, NJ tree and PC analysis revealed that all subpopulations contained both pure (non-hybrid) as well as admixed (share SNPs from different subpopulation) genotypes. For broadening genetic base of population, pure genotypes should be crossed. But for improving or introgression of specific traits, admixed genotypes could also be crossed which will reduce the population size required for phenotypic screening. However, population diversity indicates the level of similarity or dissimilarity based on alleles that not necessarily come from the same parent or ancestor, this tends to inflate the real differentiation between any two pair of individuals (96). Since a breeder would like to combine positive alleles that historically never have been combined, IBS values are good to decide what individuals will be crossed. Low IBS is the best. But, IBS values in self-pollinated crops tends to be higher than that in cross-pollinated crops as heterozygosity reduces the probability of two alleles at a locus of being identical by state (97). In our study, most of the genotypes had weak relatedness as approximately 64% of pairwise coancestry ranged from 1.21 to 1.50. Crossing among genotypes from subpopulation P2 will demonstrate more diversity, than that of other subpopulations, as most genotypic combinations of P2 shows low IBS coefficients than others. This finding is in line with the evolutionary history of B. napus where semi-winter type is the base population containing more divergence. Gradually this diversity is narrowed down in P3 (spring type, mixed origin) and P1 (winter type), because genotypic pairs belong to P3 and P1 having high IBS values evolved from semi-winter type (81). Subpopulation P4 exhibited highest number of pairs having IBS > 1.5, which is obvious as these genotypes are advanced breeding lines developed from crossing of same set of parents in different combinations. Genotypic pairs of P5 (rutabaga type) also showed high coancestry which is may be due to the duplicates. We could discard the duplicates during the crossing program.

### Linkage Disequilibrium

Linkage disequilibrium can be defined as the correlation among polymorphisms in a given population (98). The strength of association mapping relies on the degree of LD between the genotyped marker and the functional variant. Linkage disequilibrium analysis provides insight into the history of both natural and artificial selection (breeding) and can give valuable guidance to breeders seeking to diversify crop gene pools (20). SNPs in strong LD are organized into haplotype blocks which can extend even up to few Mb based on the species and the population used. Genetic variation across the genome is defined by these haplotype blocks. Haplotypes which are subpopulation-specific are defined by various demographic parameters like population structure, domestication, selection in combination with mutation and recombination events. Conserved haplotype structure can then be used for the identification and characterization of functionally important genomic regions during evolution and/or selection (99). Also, the extent of LD needs to be quantified across the genome at high resolution (down to approximately 1 Kbp) (100). The information is important for choosing crossing schemes, association studies and germplasm preservation strategies (101–104).

We used markers from across the genome to quantify the LD for the core collection. Low level of LD was evident for each individual subpopulation in A, C and whole gnome. The low level of LD can be due to multiple factors. First of all, canola is a partially outcrossing species with an average of 21-30 % of cross pollination (105–107). The outcrossing occurring in canola leads to more recombination and to a breakdown of haplotype blocks. Secondly, the ancestral history of canola is limited in comparison with other crops, such as rice, common bean, wheat and corn, restricting the selection of desirable haplotypes during the evolution. In other words, there was no adaptation or domestication pressure on the species, which would lead towards positive selection. Third, the only selection pressure imposed on the species for a relatively short time was breeding. However, the breeding practices were biased towards selection of only few phenotypes. Additionally, the short period of time under selection pressure might have not been sufficient to select favorable haplotypes in the genome. Fourth, since canola cultivars with different growth habits are compatible there has been always gene flow present between them contributing to the low level of LD. The *N_m_* >1 was observed in this study, which supports this gene flow. Finally, the restriction enzyme used to develop the libraries for sequencing of the core collection helped in identification of SNPs largely residing in genic regions, which are prone to high recombination, contributing to the low level of LD.

In this study, we have identified that the LD decay in B. napus varied across chromosomes of both A and C genomes. In addition, LD in C genome decayed much slower than A genome. C genome also contained larger haplotype blocks than A genome. This LD patterns are consistent with previous findings (20, 27, 108–110). The slower LD decay and presence of long haplotype blocks in C genome indicates that high level of gene conservation could have resulted from limited natural recombination or could be exchanged of large chromosomal segment during evolution. In the whole genome, presence of subpopulation specific haplotype blocks suggests that these regions had been experienced selection pressure for specific geographic regions adaptation. In all subpopulations, presence of shorter haplotype blocks in A genome than C genome reveals that B. rapa progenitor of B. napus containing A genome, which has been used as oilseed crop and probably being used in hybridization process. Sharing haplotype blocks by different subpopulations specially in C genome also confirms its conserved nature. The low level of LD or haplotype blocks has implications for association mapping and a proper experimentation design is necessary for utilizing a reduced set of markers by tagging major haplotypes (111). Though low LD requires more markers to pinpoint the location of various QTL, but once a marker is found to be significantly associated with a phenotype, there might be a higher probability of identifying the casual gene.

## Conclusions

This study provided a new insight to select the best parents in crossing plan to maximize genetic gain in the population. The population structure analysis showed a clear geographic and growth habit related clustering. Low LD values indicate that our collection is a valuable resource for prospect association mapping endeavors. The genetic diversity of the core collection of *B. napus* was low. Breeding efforts will need to address this issue in order to generate future hardy and high yielding varieties with resistance to many abiotic and biotic stresses. The rutabaga type showed the highest genetic divergence with spring and winter types accessions. Therefore, the breeding strategies to increase the genetic diversity may include generating population from rutabaga and spring crosses, or using rutabaga and winter crosses.

## Acknowledgements

The authors thank Mr. Andrew Ross (Department of Plant Sciences, NDSU, Fargo, ND) for his help in planting and maintaining the plants in the greenhouse and in the field.

## Supporting information

**S1 Table. List of the genotypes analyzed in this study.**

**S2 Table. Marker diversity parameters.**

**S3 Table. Subpopulation-wise marker diversity parameters.**

**S4 Table. (a) Percentage of variation explained by the first 3 axes, (b) Eigen values by axis and sample eigen vectors.**

**S5 Table. Kinship matrix.**

**S6 Table. Mean LD values according to distance.**

**S7 Table. Subpopulation-wise and chromosome-wise LD decay rate (Kb) within each subpopulation.**

**S8 Table. Subpopulation specific and common haplotype blocks.**

**S1 Fig. Histogram of IBS coefficients.**

**S2 Fig. Chromosome-wise LD decay rate (Kb) in A genome considering whole collection.**

**S3 Fig. Chromosome-wise LD decay rate (Kb) in C genome considering whole collection.**

